# Knowledge of effort modulates visual memory biases for body postures

**DOI:** 10.1101/2025.01.24.634717

**Authors:** Qiu Han, Marco Gandolfo, Jules van Dommelen, San Schoenmacker, Klemens Drobnicki, Marius V. Peelen

**Affiliations:** Donders Institute for Brain, Cognition and Behaviour, Radboud University, 6525 HR Nijmegen, the Netherlands

## Abstract

The visual memory of others’ postures has been proposed to be shaped by knowledge and expectations. For example, the visual memory of a lifted arm was recently shown to be biased downward, suggesting that observers predicted the upcoming state of the arm based on knowledge of the effort required to hold the arm up against gravity. Alternatively, the downward bias for body postures could reflect an automatic normalization towards the most frequently observed arm position, with arms more often observed in a low position. Here, in three experiments, we provide evidence that the downward bias is flexibly modulated by knowledge of effort. In Experiment 1, we found a stronger downward bias for arm postures that are relatively effortful (lifting an arm above the shoulders while standing) compared to arm postures that are less effortful (lifting an arm above the chest while lying down). In Experiment 2, we found a stronger downward bias when the actor was standing (viewed from the side) than when the actor was lying down (viewed from above), even though the arm postures were visually identical. Moreover, dividing attention during the encoding stage reduced the bias, showing that attentive processing of the stimulus was required for the bias to emerge. Finally, in Experiment 3, we found that executing the observed posture during the visual memory task did not further increase the downward bias. Together, these findings demonstrate a high-level cognitive influence on the visual memory for body postures.

## Introduction

Our internal representation of the visual environment is not a direct copy of the external world but is shaped by our knowledge, experience, and expectations (de Lange et al., 2018; Kaiser et al., 2019; Peterson, 2019; von Helmholtz, 1867). According to Bayesian theories of perception, perception follows from the integration of visual input and prior knowledge (Geisler & Kersten, 2002; Lee & Mumford, 2003). This Bayesian framework has been used to account for various perceptual and memory biases in object processing, including the finding that familiar objects appear sharper than unfamiliar ones (Perez et al., 2020) and the finding that the sizes of common objects are (mis)remembered as closer to their prototypical sizes (Hemmer & Steyvers, 2009).

One object category for which humans have extensive knowledge is the human body. Bodies are detected more easily than other objects (Gandolfo & Peelen, 2024; Stein et al., 2012) and their perception is supported by specialized brain areas in visual cortex (Peelen & Downing, 2007), effects that likely reflect our extensive visual experience with bodies (Chan et al., 2010; Stein et al., 2016). In addition to attending others’ bodies, we also gain experience about the human body by acting with it ourselves, which may shape how bodies are visually represented (C. Reed et al., 2004). Previous studies have shown that the knowledge of body biomechanics can bias the perception and memory of body movements. For example, an awkward hand rotation takes longer to imagine than an easy rotation (Parsons, 1987). Furthermore, when seeing ambiguous apparent body movements, observers tend to see the anatomically possible motion path instead of the shortest motion path (Shiffrar & Freyd, 1990, 1993). Finally, the visual memory of the end point of a movement path is usually biased along the implied motion direction (an effect called representational momentum) (Freyd & Finke, 1984; Hubbard, 2005), but this bias is reduced when the movement reaches anatomical constraints (Vandenberghe & Vannuscorps, 2023; Wilson et al., 2010).

Recently, we showed that body knowledge also affects the visual memory of *static* human body postures (Han et al., 2024): A lifted arm was remembered as lower than its actual position, presumably reflecting knowledge of gravity. Moreover, this downward bias was reduced when the arm was lifted behind the shoulder, presumably reflecting knowledge of biomechanical constraints. These results were reliably observed across experiments and tasks, but their interpretation is still unclear. Here, we considered two plausible explanations.

First, according to a normalization account, the body posture biases may reflect the statistical distribution of body postures visually experienced in the past; arms are more frequently in a lower position than high up in daily life, including during standing, walking, and sitting, which should also reflect in observers’ prior distribution of postures. Therefore, if the visual memory of an arm is drawn to the most frequent arm positions, this will manifest as a downward bias for lifted-up arms, consistent with previous results. In a similar vein, prior statistics of visual information have been shown to contribute to perception, accounting for various perceptual and memorial biases and illusions (Geisler & Kersten, 2002; Gregory, 1997). These biases most often manifest automatically, irrespectively of attention, as an attraction to the mean or median of the prior distribution. For example, overestimation of the size of small objects and underestimation of large objects (Hollingworth, 1910), motion illusions caused by a low-speed prior (Weiss et al., 2002), and attraction to the immediate recent percept (Cicchini et al., 2024; Fischer & Whitney, 2014). A similar mechanism of attraction to the mean could potentially explain the downward bias without the need to assume knowledge of gravity.

Alternatively, according to a cognitive account, the downward bias may reflect a more flexible, cognitive influence, reliant on a conceptual understanding of gravity. We know that holding an arm up requires active muscle strength and is thus effortful. In the absence of such strength, the arm will most likely be pushed down by gravity. Based on this knowledge, participants might have predicted the future state of the observed figure which led to a biased memory. Evidence for high-level effects on memory biases comes from a study showing that knowledge of an object’s typical motion (e.g., knowing that a rocket can move) modulates the strength of representational momentum (Reed & Vinson, 1996; Vinson & Reed, 2002). In the domain of action perception, studies also showed that the perception of other’s actions is shifted forward and that this effect incorporates knowledge of the actor’s intention (Hudson, Nicholson, Ellis, et al., 2016; Hudson et al., 2018).

The current series of experiments was designed to tease apart these accounts by manipulating variables that change the observer’s conceptual understanding of the effect of gravity on body posture. If the downward body bias follows a higher-level understanding of effort, we would expect that the bias can be modulated by contextual variables that influence the effort needed to maintain the observed posture (e.g., a reduced bias when viewing a person lying down). By contrast, if the downward body bias is an automatic effect of prior visual experience, we would not expect contextual variables to influence the bias. In Experiment 1, we compared the downward bias for postures that are known to require more or less effort to maintain. In Experiment 2, we presented bodies with lifted arms in a standing position, requiring effort, or lying on a bed (viewed from above), not requiring effort. Finally, in Experiment 3, we manipulated the observer’s own posture to test whether activating knowledge of effort in this way would modulate the downward memory bias.

## Experiment 1

### Introduction

To test whether the downward bias is modulated by knowledge of effort, in Experiment 1 we compared the downward bias for a posture that is relatively effortful to maintain with a posture that is easier to maintain. In the Standing posture, the actor’s arm was positioned above the shoulder, while in the Lying posture, the arm was positioned above the chest (Fig 1A). The elbow angle and orientation were identical in both conditions. However, holding the arm at that angle while lying down requires less effort since the pectoral muscles recruited when lying down are stronger than the shoulder muscles recruited when standing up.

**Figure 1:**
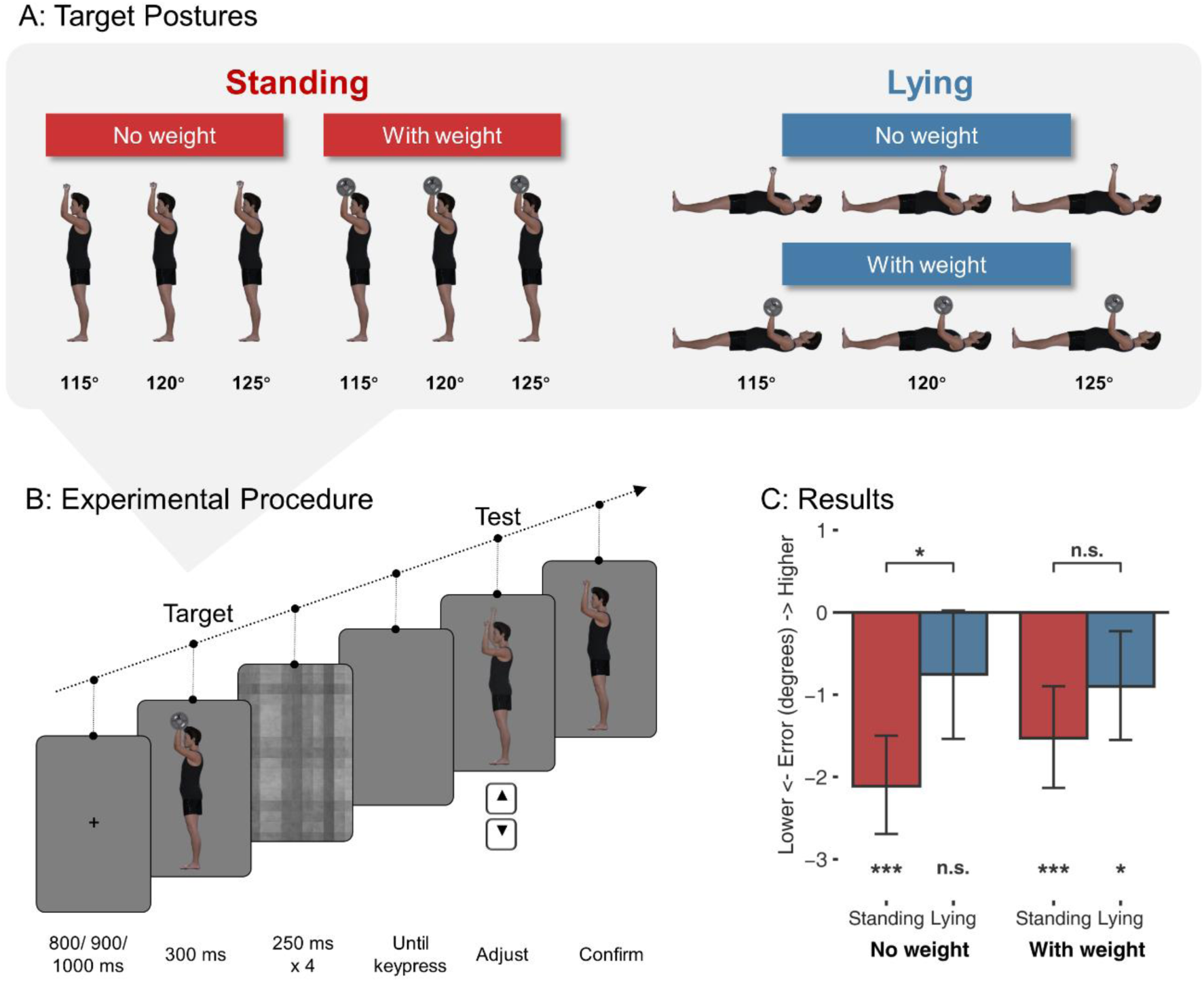
Stimuli, procedure, and results of Experiment 1. A: All target postures used in the experiment. B: Schematic of trial structure. After the fixation and the target posture (here the standing-with weight condition as an example), four dynamic masks appeared, after which the participants could press a key to start adjusting the arm to match the target posture. The arm angle on the screen dynamically changed with the keypress of the up/down arrow (illustrated by the transparent arm in the figure which was not shown during the experiment). C: Results of Experiment 1. The graph shows the mean adjustment error for the four conditions. Errors below 0 reflect a downward bias, such that participants misremembered the posture to be lower than it was. Error bars show 95% CI; ***: p < 0.001, **: p < 0.01, *: p < 0.05, n.s.: not significant.

As a further manipulation of effort, we manipulated whether the actor was holding a bar with or without weight (Fig. 1A). If the downward bias is modulated by knowledge of effort, the downward bias should be larger for Standing than Lying postures, and larger for With-weight than No-weight conditions.

### Method

#### Participants

Using a power analysis (Jamovi) with 80% power for detecting a medium effect size (d = .5) assuming a two-sided criterion for detection, with a maximum type I error rate of α = .05, we predetermined a desired sample size of 34. In consideration of potential exclusions due to poor task performance, we recruited N = 37 participants (age M = 30.51, range = [19, 68], 23 females, 14 males) who took part in the study and gave informed consent for participation in the experiment. The participants were recruited through the participant panel of the university (SONA) in return for course credits or by voluntary participation. The study was approved by the ethics committee of Radboud University (reference number ECSW-LT-2024-4-11-64796).

#### Stimuli

Body posture images were generated by rendering human figures in DAZ Studio 4.22 (Daz Productions, Inc). The target stimuli depicted a profile view of a male figure with a weight-lifting posture, either standing or lying down, shown holding a bar with or without extra weight (Fig. 1A). For each lifting posture, three slightly different angles were used to introduce some variation for the target stimuli: the upper arms (shoulder to elbow) were angled at 115, 120, or 125 degrees relative to the direction of gravity; that is, the larger the angle is, the higher the arm is, while the forearm was always upright. To avoid potential confounds caused by visual field differences, both left- and right-side views of the figure were generated through horizontal flipping. In total, each combination of Posture and Weight contained six images (three angles x two views). The images were presented at 533 x 300 pixels for the lying posture and 300 x 533 pixels for the upright posture. The mask image was a monochrome checkerboard pattern extending 500 by 500 pixels.

#### Procedures

The experimental procedure was programmed using the JsPsych library (de Leeuw, 2015) and the psychophysics plugin (Kuroki, 2021). The experiment was conducted on a laboratory monitor with a resolution of 1920 by 1080 pixels and physical dimensions of 527 mm x 296 mm. Participants were given a verbal as well as a written explanation of the trial procedure. Two practice rounds of five trials each were given before the experiment started.

At the beginning of each trial, a fixation cross appeared at the center of the screen for 800, 900, or 1000 ms (Fig. 1B). Then, the target posture was shown for 300 ms, followed by four consecutive checkerboard mask images for 250 ms each to eliminate potential aftereffects. After the mask images, the participants were instructed to press an initiation key to start the adjustment phase. This initiation key was either a left or right arrow key, counterbalanced across participants. Upon pressing this key, the same posture would show up again, but with an angle randomly sampled from a range of 90 to 140 degrees. During the adjustment phase, the human figure held nothing in the hand to avoid the use of the weight’s position as a reference and to avoid the weight-lifting movement appearing effortless. Participants could then use the up and down arrow keys to adjust the angle of the figure’s arm at a step size of 2.5 degrees to match the target posture as closely as possible. They could keep adjusting until the posture matched their remembered posture, then confirm the position they decided on by pressing the space bar. Participants were given a maximum of 10 seconds for the adjustment phase after the mask offset. A warning message was given when the participant skipped the adjustment phase by directly pressing the space before pressing the initiation key or when they took too long (> 3 seconds) to initiate the adjustment phase.

Each participant took part in all conditions. For each Posture (Standing, Lying) x Weight (No weight, With weight) combination there were 24 trials (three angles x two side views x four repetitions), resulting in a total of 96 experimental trials. The trial order was fully randomized. The trials were divided into four blocks, with short breaks in between. At the end of each block, participants received feedback on their average absolute error to maintain engagement and motivation. At the end of all trials, participants were debriefed.

#### Analysis

One participant was excluded due to a misunderstanding of the task instructions. Trials in which the adjusting image was not initiated within 3 seconds after the mask offset and trials in which the participants skipped the adjustment were discarded. Trials with an absolute error larger than 15 degrees were also discarded, as these likely indicated a lapse of attention (Han et al., 2024). Taken together, 3.27% of trials were excluded from further analysis.

For the remaining trials, the error was calculated by subtracting the presented angle from the adjusted angle, so that a negative error means that the adjusted angle was lower than the target angle. Posture (Standing, Lying) and Weight (No weight, Wiith weight) are the two independent factors. Trials belonging to the same combination of conditions were averaged to get the dependent measure of mean error. A two-way repeated measures ANOVA with Posture and Weight as within-subject factors and mean error as the dependent variable was conducted. A simple effects analysis was then used to determine the effect of Posture at each level of Weight. The assumptions of normality and equal variance were checked. Additionally, a t-test against zero was conducted for each condition to test for the presence of a downward bias.

The data were preprocessed and analyzed using the Python libraries pandas (The pandas development team, 2010) and pingouin (Vallat, 2018). The data were visualized using the Python library seaborn (Waskom, 2021).

### Results

As shown in Fig. 1C, in all conditions except for the Lying-No weight condition, the error was significantly below zero, indicating a downward bias (Lying-No weight: M = -0.75, 95% CI [-1.54, 0.03], t(35) = -1.95, p = .059; Lying-With weight: M = -0.90, 95% CI [-1.60, -0.20], t(35) = -2.61, p = .013, d = 0.43; Standing-No weight: M = -2.11, 95% CI [-2.79, -1.44], t(35) = -6.35, p < .001, d = 1.06; Standing-With weight: M = -1.53, 95% CI [-2.19, -0.87], t(35) = -4.69, p < .001, d = 0.78).

The two-way repeated-measures ANOVA showed no main effect of Posture (F(1,35) = 3.46, p = .071, η²_p_ = .09) and no main effect of Weight (F(1,35) = 2.32, p = .137, η²_p_ = .06). The interaction between Posture and Weight was significant (F(1,35) = 4.73, p = .037, η²_p_ = .12). A simple effects analysis revealed that the downward bias was larger for the Standing than the Lying postures within the No-weight condition (t(35) = 2.38, p = .023, d = .63), but not within the With-weight condition (t(35) = 1.14, p= .260, d = .31; Fig. 1C).

### Discussion

The results of Experiment 1 showed a larger downward bias for the more effortful (Standing) posture compared to the less effortful (Lying) posture when there was no extra weight, suggesting that the downward bias was modulated by the knowledge of effort. However, in the presence of extra weight, this difference did not reach significance. Furthermore, contrary to our expectation, the added weight did not generally increase the downward bias. A possible explanation for this finding could be that the added weight led some participants to consider the postures in the context of a weight lifting exercise, for example as a bench press or a push press exercise. If so, those participants may have expected the arm to go up instead of down.

Another possible explanation for the lack of a weight effect is that the weight attracted attention during the encoding phase, reducing attention to the body (which was also partly occluded). Furthermore, the weight was removed in the adjustment phase to avoid it being used as a reference, causing a mismatch of visual features between the memory encoding and retrieval stages. Therefore, the No-weight condition was better controlled in terms of visual features, because no visual differences were introduced between the target and to-be adjusted posture. Taken together, although the No-weight condition provided evidence that knowledge of effort modulates the downward bias, the with-weight condition remains inconclusive. In Experiment 2, we manipulated effort differently, now keeping the upper-body postures the same across conditions but changing the context around the body.

## Experiment 2

### Introduction

Experiment 2 tested whether the downward bias for body postures is modulated by knowledge of effort by manipulating postures in a different way: The same figure lifting her arm was shown either standing seen from the side or lying on a bed seen from above (Fig. 2B). The standing posture requires effort to prevent the arm from being pulled down by gravity, while the lying posture does not. Compared to Experiment 1, the current manipulation better controlled the potential confound of visual feature differences: due to the identical visual features of the upper body between Standing and Lying conditions, the effort information can be extracted only based on the context around the human figure.

**Figure 2:**
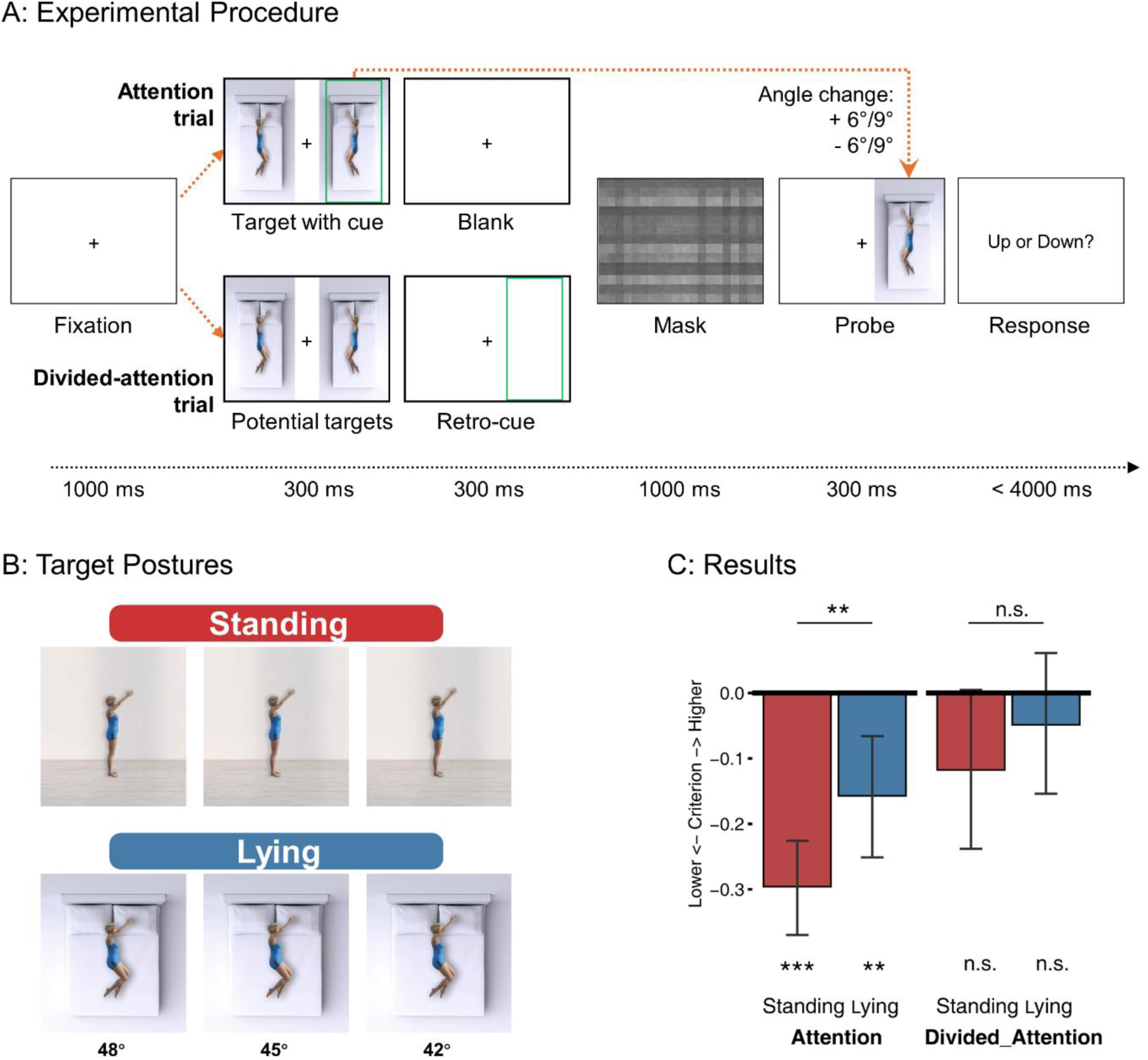
Stimuli, procedure, and results of Experiment 2. A: Schematic of trial structure, with the lying condition as an example. In the Attention condition, the cue appeared simultaneously with the target; in the Divided attention condition, the retro-cue appeared after the two potential targets disappeared. For both conditions, the probe appeared on the same side of the target after a checkerboard mask. Participants indicated whether the arm in the probe moved up or down compared to the remembered target. B: All target postures used in the experiment. During the experiment, the images were squeezed horizontally to fit on the screen as shown in A. C: Results of Experiment 2. Bars show criteria of four conditions, with negative criteria corresponding to a tendency to judge the probe arm more often as higher than the target, indicating a downward memory bias. Results showed a main effect of Posture, with a larger downward bias for Standing than Lying conditions, and a main effect of Attention, with a larger downward bias for Attention than Divided attention conditions. Error bars show 95% CI; ***: p < 0.001, **: p < 0.01, *: p < 0.05, n.s.: not significant.

To further probe the cognitive nature of the downward bias, we also tested whether the downward bias is modulated by attention in Experiment 2. If the downward bias reflects an automatic perceptual effect, it may not depend on the degree of attention allocated to the stimulus, similar to other automatic perceptual processes (e.g., grouping; Lamy et al., 2006; Rashal et al., 2017). By contrast, if the downward bias relies on higher-level cognitive processing, it will require attentional resources and thus depend on the degree of attention allocated. Here, we manipulated attention using a cueing paradigm to direct attention to either a single human figure (Attention condition) or two human figures (Divided-attention condition) at the posture encoding stage (Fig. 2A). In the Divided-attention condition, the relevant figure was cued after the stimuli had already disappeared, requiring participants to divide attention across the two stimuli during encoding. To easily implement the attentional manipulation, and to avoid a possibly unnatural adjustment of the posture while lying down, in this experiment we used a posture change discrimination task (Han et al., 2024) instead of a self-paced posture adjustment task. In this task, participants judge whether a lifted arm appears up or down compared to previously observed target postures (Figure 2A).

### Method

#### Participants

This experiment included the same 37 participants as in Experiment 1. The order of experiments was counterbalanced across participants. Two participants did not finish this task due to technical issues, resulting in a final sample size of 35.

#### Stimuli

Body images were created with DAZ studio 4.22 (Daz Productions, Inc). A standing female figure was shown from the side, lifting her arm to her front. The background was either a wall seen from the side (the Standing condition) or a bed seen from above (the Lying condition). The upper body of the figure was identical in the two conditions, but the leg in the Lying condition was modified to depict a realistic lying posture (Fig. 2B). Different from Experiment 1, the vertical upwards posture was coded as the zero point, such that larger angles correspond to a lower position.

The body images were 214 pixels wide and 380 pixels high. The cue was a green frame larger than the body’s size. The masking image was a grey-scale checkerboard image similar to Experiment 1.

#### Procedures

The experimental procedure was created using the PsychoPy Builder (v2023.2.3). Experiment 2 used a version of the task (change discrimination) in which the participant does not directly adjust the arm but judges whether the arm position of the probe posture is higher or lower compared to an earlier presented target. If the target’s arm is remembered as lower than it actually was, an upward change in the probe posture should be more noticeable. Thus, an asymmetry in detecting upward and downward changes indicates a bias in visual memory. This task was previously shown to result in similar downward biases as the adjustment task used in Experiment 1 (Han et al., 2024). The order of trials corresponding to the two attention conditions (Attention, Divided-attention) was randomized across all blocks.

Each trial started with the presentation of a fixation cross at the center of the screen for 1000 ms, followed by two postures of the same condition (i.e., both Lying or both Standing) but with different angles. The two figures were facing each other, one at either side of the fixation cross (Fig. 2A). One of the two images was the target to be remembered and compared to a later probe image. For Attention trials, but not for Divided-attention trials, one of the two images was shown together with a green frame signaling the target. For Divided-attention trials, a retro-cue (300 ms) was shown only after the two images disappeared, such that participants had to divide attention across the two images. To equate the memory maintenance periods of the two trial types, a blank screen was shown in the Attention condition after the target had disappeared for the same period as the retro-cue (300 ms) (Fig. 2A). The target posture’s angle could be 42, 45, or 48 degrees, and the non-target posture’s angle was 39, 42, 45, 48, or 51 degrees.

After a mask of 1000 ms, a probe stimulus was shown for 300 ms (Fig. 2A). The probe’s arm had moved up or down by 6 or 9 degrees relative to the target. Participants were required to indicate whether the arm of the probe had moved up or down compared to the cued target by pressing the “up” or “down” arrow key. The text ‘Did it move up or down?’ was displayed after the probe offset for a maximum of four seconds or until either key was pressed. Participants were instructed to respond as quickly and accurately as possible and keep their eyes on the fixation cross at all times during the trial.

Each factor level combination (Attention – Standing, Attention – Lying, Divided attention – Standing, Divided attention – Lying) consisted of 48 trials. The experiment consisted of 192 trials in total, divided over 4 blocks with breaks in between. While Attention and Divided-attention trials were fully randomized, the Posture factor was blocked to ensure unambiguous understanding of the context, meaning that each block consisted solely of either the Standing or Lying condition. To control for order and learning effects, the order of blocks was counterbalanced across participants, with every other participant receiving either the ABAB or BABA order. Before the experiment, a practice block composed of three trials of each of the four types of conditions was provided for participants to get acquainted with the task.

#### Analysis

The downward bias was quantified by using criterion (c) from signal detection theory (c = – (z(hit) + z(false alarm)) / 2). We arbitrarily considered upward change as the signal; thus, a negative criterion indicates a bias towards choosing upward change over downward change, suggesting that changes in the upward direction were more noticeable to the participants. Criteria for all participants per condition were tested against zero using two-tailed t-tests. The effects of the two experimental factors were tested using a two-way repeated measures ANOVA. Data analysis was conducted in R using BruceR package (Bao, H.-W.-S., 2021).

### Results

As shown in Fig. 2C, both postures showed a significant downward bias in the Attention condition (Standing: M = -0.27, 95% CI [-0.37, -0.22], t(34) = -7.70, p < .001, d = 1.30; Lying: M = -0.16, 95% CI [- 0.25, -0.06], t(34) = -3.34, p = .002, d = 0.57) but not in the Divided-attention condition (Standing: M = -0.12, 95% CI [-0.24, -0.001], t(34) = -2.02, p = .051, d = 0.34; Lying: M = -0.049, 95% CI [-0.16, -0.06], t(34) = -0.90, p = .37, d = 0.15).

A repeated measures ANOVA revealed a significant main effect of Posture (F(1, 34) = 7.00, p = .012, η²_p_ = 0.17), indicating that the arm in the standing context was remembered as lower than the arm in the lying context. The main effect of Attention (F(1, 34) = 20.6, p < .001, η²_p_ = 0.38) showed that the downward bias was larger when attention was focused on the target stimulus. The interaction between Posture and Attention was not significant (F(1, 34) = 0.97, p = .33, η²_p_ = 0.028).

### Discussion

Experiment 2 provides further support for a higher-level cognitive influence on the downward bias for body postures. Using context to manipulate the effort required for maintaining a posture, we again revealed that more effortful postures lead to a larger downward bias. In this experiment, the task-relevant part of the body (the upper body) was visually identical for the standing and lying conditions. Therefore, if the bias reflects an automatic attraction to the mean arm position, we should expect no difference between these two conditions. Yet, when the context implied that the figure was lying on a bed, so that the lifted arm was not pushed down in the direction of gravity, the downward bias was reduced. This suggests that contextual information was incorporated by the viewer when forming an understanding of the effort required by the actor to hold the lifted arm posture, modulating the downward bias. These findings demonstrate that the downward bias for body postures is, at least partly, caused by knowledge of the effort to maintain a posture against gravity.

A second finding of Experiment 2 was that the downward bias was reduced when less attention was allocated to the body stimulus. This finding suggests that the downward bias requires controlled, attentional processing at the encoding stage, corroborating the idea of a cognitive contribution to the downward bias for body postures.

## Experiment 3

### Introduction

Experiments 1 and 2 both revealed a cognitive modulation of the downward bias for body postures, supporting the hypothesis that this bias was based on the knowledge of effort caused by gravity instead of (or in addition to) normalization to the mean observed arm position. In these experiments, participants extracted information about effort from the visual stimuli, that is, by inferring if the figures were lifting their arms against gravity given the visual context. In Experiment 3, we manipulated the cognitive factor not by manipulating the stimuli but by manipulating the participants’ own motor state (Barsalou, 2008). Although postures and actions can be understood by visual analysis (Allison et al., 2000; Lingnau & Downing, 2015; Peelen & Downing, 2005; Vannuscorps & Caramazza, 2016b), studies have also found that the observer’s own actions can modulate this process (Galvez-Pol et al., 2020; Hamilton et al., 2006). Therefore, we tested whether the understanding of other people’s efforts can be modulated by the effort that observers experience themselves: executing the same arm-lifting posture as the one observed might activate the knowledge of the effort required to hold up the arm. The direct sensorimotor experience of the effort, on top of the knowledge extracted via the visual analysis of the stimulus, may increase the downward bias.

In Experiment 3, participants were asked to lift their arms while observing the postures on the screen. If the understanding of others’ effort is modulated by one’s own motor state, we expect that lifting an arm will increase the downward bias. By contrast, if visual processing is sufficient for extracting the effort information, we expect no effect on the executed posture. Furthermore, in this experiment, we tested two different arm-lifting postures: in front of the body (Front) and behind the body (Back). We included back postures because based on our previous work (Han et al., 2024), memory for these postures is also biased by biomechanical constraints besides by gravity (an arm lifted behind the back, can’t go further back, due to the body structure, see Fig. 3A). Such body-specific constraints may influence the visual memory of the posture via participants’ sensorimotor knowledge of how effortful they are, leading to increased biomechanical biases during execution, as compared to a rest condition.

**Figure 3.**
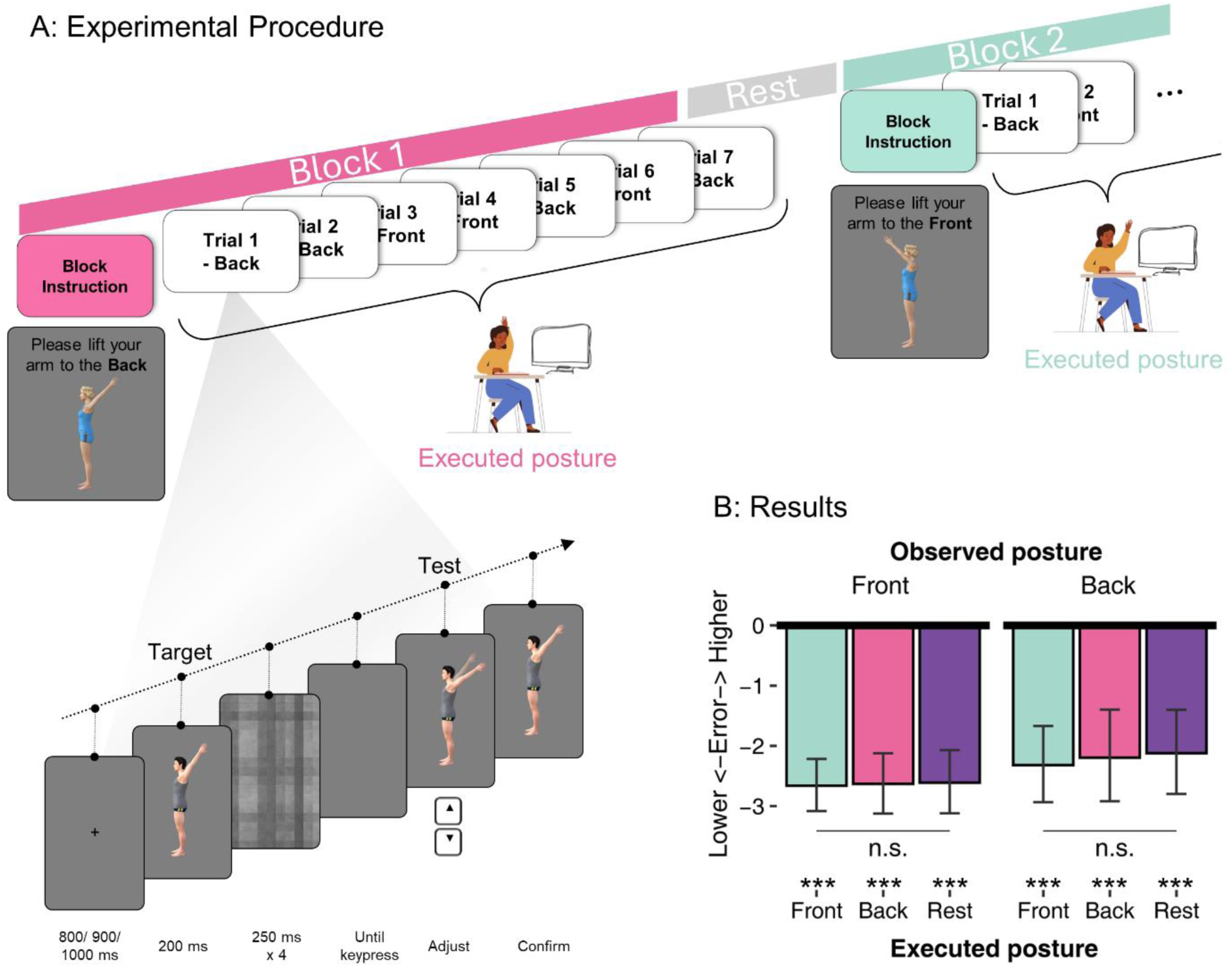
Procedure and results of Experiment 3. A: Experimental procedure. Each block started with an instruction about the executed posture in this block: Lift to the front, lift to the back, or rest. Participants needed to execute this posture with their left arm while conducting seven trials with their right hand. The Observed postures (Front or Back) in one block were randomized. After each block, a mandatory rest was given, and then the next block started with the instruction of the next executed posture. On the bottom left is the structure of an example trial, here showing a Back posture of a male figure facing left. The task is similar to that illustrated in Fig. 1B. B: Results of Experiment 3. All conditions showed a downward bias. Executed posture did not influence the downward bias for either observed posture. Error bars show 95% CI; ***: p < 0.001; n.s.: not significant.

### Method

#### Participants

Participants were recruited through the SONA platform in exchange for course credits. 38 different participants were recruited based on the predetermined sample size of 34 (see Experiment 1). One person’s demographic data were missing. For the other 37 people: age mean = 21.27, range = [18, 32]; 28 females, 9 males. All participants gave written consent.

#### Stimuli

Posture images were identical to those used in previous work (Han et al., 2024): rendered male and female figures lifting their arms to the front or the back of their body using DAZ studio 4.15 (Daz Productions, Inc). For each posture, seven slightly different angles were used to introduce some variation: Taking the vertical up as angle 0, the arm could be at an angle of 36, 39, 42, 45, 48, 51, or 54. Both left and right views were used. Body images were 300 pixels wide and 480 pixels high. Mask images were 350 pixels wide and 525 pixels high. Masks were presented slightly larger than the body to achieve a better masking effect.

#### Procedures

During the experiment, participants executed an arm posture with their left hand while conducting the task using their right hand (Fig. 3A). To avoid fatigue, trials were grouped into short blocks of seven trials of the same Executed posture. Before each block, a prompt indicated which posture the participant had to execute with their left arm: lifting to the front, lifting to the back, or rest. A rendered figure executing the same front or back posture at 45 degrees was shown together with the prompt (Fig. 3A); participants were instructed to imitate the posture as closely as possible without experiencing discomfort. Participants were instructed to start the block only after they had put their left arm in the corresponding posture and to hold this posture until finishing the seven trials of that block. The task was the same as the adjustment task in Experiment 1 except for some minor differences: since the postures here were simpler in terms of visual features, the target image was presented for 200 ms (Fig. 3A), and the adjustment range was from 30 to 60 degrees (corresponding to 120 to 150 degrees in Experiment 1). After each block, participants were encouraged to rest as long as they needed, especially if their arm was tired.

Before the experiment, two practice blocks of six trials each were given to familiarize participants with the task. After each practice block, participants were provided with their average absolute error during that block as feedback. During the experiment, each executed posture block was repeated eight times in random order, resulting in 24 blocks (168 trials) in total. Within a block, the figure (male, female), the stimulus posture (Front or Back), and the specific angle were all randomized, while the facing direction (left, right) was kept the same.

#### Analysis

Exclusion followed the same rules as Experiment 1, resulting in 3.21% of trials being excluded overall. No participants had to be excluded. To allow inference on null effects, we calculated the Bayes Factor (BF) for a two-way ANOVA with the Executed posture and the Observed posture as within-subject factors using the BayesFactor R package. We reported BF_01_ (the ratio of the posterior probability of the null hypothesis against the alternative hypothesis), for which a BF_01_ larger than 3 conventionally indicates some evidence supporting the null hypothesis, a BF_01_ larger than 10 indicates strong evidence, while a BF_01_ < 0.3 or 0.1 suggest some or strong evidence for the alternative hypothesis (Schmalz et al., 2021). A frequentist two-way ANOVA was also conducted where Greenhouse-Geisser was used to correct for sphericity violations.

### Results

As shown in Fig. 3B, all six conditions showed a gravity bias (ps < 1.08E-6, ds > 0.94, BF_01_s < 6.29E-5). The frequentist ANOVA did not show significant main effects or interaction (Executed posture: F(1.56, 57.8) = 0.36, p = .65, Observed posture: F(1, 37) = 1.91, p = .18, interaction: F(1.85, 68.5) = 0.19, p = .81), suggesting a lack of effect of one’s executed posture on the bias. Consistently, Bayes Factors suggested strong evidence supporting a null effect of Executed posture (BF_01_ = 18.5), and no interaction between Executed posture and Observed posture (BF_01_ = 11.6). There was some evidence for an effect of Observed posture (BF_01_ = 0.36), with a smaller downward bias for back postures than front postures (Fig. 3B), consistent with knowledge of biomechanical constraints that prevent the back posture from going down further (Han et al., 2024). Executed postures did not affect the observation of the front posture (BF_01_ = 12.0) nor the back posture (BF_01_ = 7.43).

### Discussion

In Experiment 3 we observed that executing an arm posture did not increase the bias of visual memory of observed postures. This result suggests that visual information may have been sufficient for activating knowledge of effort. However, we cannot exclude that under different circumstances, concurrent action execution influences visual perception and/or visual memory. One possible reason for the lack of a modulatory effect of action execution is that this effect might have been masked by the additional task demand (relative to the Rest condition) required for executing the postures. As shown in Experiment 2, when attention is drawn away from the visual stimuli, the visual memory bias is also reduced, which might counteract the effect of action execution.

The contribution of motor information to visual processing has been a long-discussed topic, especially for the visual processing of actions and postures. Some studies have argued for a causal role of motor simulation in action and posture understanding (Zentgraf et al., 2011), supported by the finding of cortical activation shared between action execution and action observation (Buccino et al., 2004; Järveläinen et al., 2004; but see Lingnau et al., 2009). These motor simulation theories may predict a facilitatory or inhibitory effect of posture execution on posture observation, which was not observed here. However, it should be noted that our experiment was not designed to provide a test of these theories; for example, our task probed visual memory rather than action understanding. Furthermore, the effect of action execution on the sensory-motor cortex might be most prominent when observing postures in a first-person perspective (Jackson et al., 2006) rather than the third-person perspective used here, and may only serve a cognitive function when mapping the goal of others onto the observers themselves (Rizzolatti & Sinigaglia, 2010).

### General Discussion

In the current series of experiments, we found that the downward visual memory bias for body postures was modulated by the observer’s knowledge of the effort required to hold the postures: more effortful postures (e.g., lifting an arm while standing) led to a larger bias than less effortful postures (e.g., lifting an arm while lying down). We also found that reduced attention during the encoding stage significantly reduced the bias, indicating that the biases required attentive processing of the stimuli. Finally, we found that observers executing the observed posture did not further increase the downward bias. Together, these findings support the hypothesis that the downward bias is flexibly modulated by higher-level inferences based on prior knowledge, rather than only reflecting an automatic normalization process towards the mean body posture. Moreover, visual information appears to be sufficient for activating this knowledge, such that executing the observed posture does not further increase the downward bias.

Understanding other people’s actions and mental states is an essential ability for human interaction (Teufel et al., 2010). In the current experiments, viewers were able to rapidly infer the effort of other people by observing their posture and combining this posture information with information about the context (e.g., bed or wall) and prior knowledge of gravity orientation and muscle strength. The resulting modulation of the downward memory bias suggests that the visual system predicts potential future body states of other people (Bach & Schenke, 2017): The more effortful the posture is, the more likely it is that the arm will fall. This prediction then biases the visual memory of the posture.

Although it is the first time that such a modulation is found for static postures with no implication of any movement, effects of prediction on memory have previously been observed for object motion and human motion (Freyd & Finke, 1984; Hubbard, 2005). These prediction-related memory biases are often detrimental to performance in the specific task setting of an experiment, which typically requires participants to be as precise as possible, but lend important benefits for living in a dynamic world: These biases situate people further along the direction in which the event will most likely unfold, thus preparing them for reactions like interception and interaction. Similar to the current findings, these biases can be modulated by contextual information, including object properties (C. L. Reed & Vinson, 1996; Vinson & Reed, 2002), object location (Bertamini, 1993), and other people’s intentions and goals (Hudson, Nicholson, Ellis, et al., 2016; Hudson, Nicholson, Simpson, et al., 2016). These phenomena, together with our findings, reveal a flexible use of context information to optimize predictions of upcoming states, serving an adaptive function in a changing environment.

The observed modulation of the downward bias by relatively high-level cognitive factors refutes the explanation that the biases arise solely from a normalization process, which should yield the same amount of bias regardless of the context. Instead, observers were able to flexibly employ different priors not only based on different postures (that lifting an arm is more difficult when standing than lying) but also based on the understanding of the whole scene (that gravity does not pull the arm downward when lying on the side). All available information is combined to ensure better prediction, in line with Bayesian accounts of perception and decision making (Bays et al., 2024; Knill & Saunders, 2003).

Experiment 3 demonstrated that lifting an arm while performing the task did not modulate the body posture biases. While there may be several reasons for such a null result (see Discussion of Experiment 3), one interpretation of this finding is that knowledge of effort does not depend on motor simulation but can be gained through cognitive inference based on visual information alone. Note, however, that this experiment does not speak to the question of whether motor experience is required for forming body-related knowledge in the first place. One possibility is that motor experience is necessary during knowledge acquisition but not during perception. Alternatively, observation of other people’s actions and postures might be enough for acquiring body-related knowledge. To test the role of motor experience in acquiring the knowledge of effort, future work can test individuals who lack specific motor experience (Vannuscorps & Caramazza, 2016a, 2016b) or who have additional motor experience such as dancers.

In summary, the current results show that visual memory biases for body postures are flexibly modulated by knowledge of effort, rather than reflecting an automatic normalization towards frequently seen postures. Observers are able to infer other people’s efforts through visual analysis of the posture embedded in its environment, and adjust their expectations accordingly. Future studies could test a larger range and variety of postures, in different contexts, to create a more comprehensive understanding of how knowledge affects visual memory of body postures.

## Acknowledgments

We thank Harry Steinharter, Berkay Serçeoğlu, Martin Wimmers, and Lydia Moonen for collecting data. The study is funded by China Scholarship Council (CSC) and European Union’s Horizon 2020 research and innovation program under the Marie Skłodowska-Curie (grant agreement No. 101033489).

## Author contributions

## Open Practices Statement

All the data and materials of all experiments are available online.

Data and scripts for Experiment 1&2: https://data.ru.nl/collections/di/dcc/DAC_2024.00072_381 Data and scripts for Experiment 3: https://data.ru.nl/collections/di/dcc/DAC_2023.00028_869

